# Full-length Cryptochrome 1 in the outer segments of the retinal blue cone photoreceptors in humans and great apes suggests a role beyond transcriptional repression

**DOI:** 10.1101/2024.10.10.617617

**Authors:** Rabea Bartölke, Christine Nießner, Katja Reinhard, Uwe Wolfrum, Sonja Meimann, Petra Bolte, Regina Feederle, Henrik Mouritsen, Karin Dedek, Leo Peichl, Michael Winklhofer

**Author notes:** To whom correspondence should be addressed: (RB), (LP), (MW). Scuola Internazionale Superiore di Studi Avanzati (SISSA), Trieste, Italy (Katja Reinhard).

## Abstract

Mammalian cryptochrome 1 (CRY1) is a central player in the circadian transcription-translation feedback loop, crucial for maintaining a roughly 24-hour rhythm. CRY1 was suggested to also function as blue-light photoreceptor in humans and has been found to be expressed at the mRNA level in various cell types of the inner retina. However, attempts to detect CRY1 at the protein level in the human retina have remained unsuccessful so far. Using various C-terminal specific antibodies recognizing full-length CRY1 protein, we consistently detected selective labelling in the outer segments of short wavelength-sensitive (SWS1, “blue”) cone photoreceptor cells across human, bonobo, and gorilla retinae. No other retinal cell types were stained, which is in contrast to what would be expected of a ubiquitous clock protein. Subcellular fractionation experiments in transfected HEK cells using a C-terminal specific antibody located full-length CRY1 in the cytosol and membrane fractions. Our findings indicate that human CRY1 has several different functions including at least one non-clock function. Our results also raise the likely possibility that several different versions of CRY1 exists in humans. We suggest that truncation of the C-terminal tail, maybe to different degrees, may affect the localization and function of human CRY1.

## Introduction

Cryptochromes (CRYs) are signal transduction proteins which share significant sequence homology with light-absorbing DNA-repair enzymes known as photolyases (1–4). Of the six major subgroups of cryptochromes that have been identified in animals (5), only two (CRY1 and CRY2) are present in mammals. Both CRY1 and CRY2 primarily function as light-independent transcriptional repressors within the circadian rhythm’s nuclear transcription-translation feedback loops ((6, 7), see (8) for review). In this feedback mechanism, the heterodimeric transcription factor CLOCK:BMAL1 binds to the regulatory E-box region of the *Period* (*PER*) and *CRY* genes and activates their transcription (9). CRY accumulates in the cytoplasm, then translocates, typically in complex with PER, back into the nucleus, and binds to CLOCK:BMAL1, thereby repressing transcriptional activation of their E-box and closing the negative limb of the feedback loop (7, 10).

While the photolyase homology region (PHR) constitutes the core structure of cryptochromes and is remarkably conserved with photolyases, mammalian CRY1 and CRY2 feature a highly flexible and diverse C-terminal tail (11). This C-terminal tail encompasses essential structural elements and functional motifs crucial for CRY1’s overall functionality, but the exact mechanisms remain elusive. In vitro experiments have indicated a role of the C-terminal tail of CRY1 in modulating circadian clock oscillations by regulating binding with CLOCK:BMAL1 (12). Additionally, a phosphorylation site within the C-terminal tail has been shown to regulate CRY1’s stability and subcellular localization, probably fine-tuning the circadian clock-machinery (13).

In addition to their established role in the transcriptional feedback loop, mammalian cryptochromes may also act as blue-light receptors in nonvisual photoreception, such as pupillary reflex and photoentrainment of circadian rhythms (14–18). With the discovery of melanopsin (19, 20), cryptochromes were no longer considered necessary for these purposes (21). However, neither in melanopsin knockout (*Opn*4^-/-^) mice (21, 22) nor in Cry1/2-double knockout mice (16) was the pupillary reflex eliminated completely, which suggests some functional redundancy among opsins and cryptochromes in the transduction of light information, which leads to behavioral modulation (15).

On the other hand, a photoreceptive function for mammalian CRYs was seriously questioned, as mammalian CRYs obtained from heterologous (bacterial and insect) expression systems (23, 24) did not bind (K_D_ > 16 µM) (24) the flavin adenine dinucleotide (FAD) co-factor that is necessary to make CRYs light-sensitive (25, 26). Nonetheless, several lines of indirect evidence argue for mammalian cryptochromes as candidate photopigments: Vanderstraeten and colleagues (27) reason that both wavelength sensitivity and the light sensitivity threshold for light entrainment of retinal biorhythms favor CRY1 and/or CRY2 as a photoreceptor over all other known light receptors in the mammalian retina. Moreover, some biochemical studies suggest that human CRYs may bind FAD *in vivo*, implying that heterologous protein expression systems may not replicate the necessary molecular requirements for FAD incorporation (23, 28–30). To understand any potential additional functions of human CRY1, it would be highly useful to know where in the human retina CRY1 is predominantly expressed, but so far immunoreactivity could not be detected in the human retina (18).

A predominantly nuclear localization would argue for a clock function, whereas the presence of CRY1 in the outer segments of shortwave-sensitive SWS1 cones in some mammalian retinae (31) and of avian CRY1a in UV/violet cone outer segments in birds (32, 33) challenge our traditional understanding of the protein’s function, as it suggests a role beyond transcriptional repression. Especially the absence of broad nucleic staining led to the suggestion that the gp-α-CRY1 antiserum labels a light-activated conformation of CRY1a with the C-terminal tail folded outward, thereby exposing the epitope for recognition by the antiserum. However, while subsequent research confirmed avian CRY1a localization in bird UV/violet cone outer segments, no difference between light- and dark-adapted retinae was observed, opposing the notion that this antiserum would selectively recognize a light-activated form (33, 34). Nevertheless, the localization of CRY1 in the outer segments far from the nucleus of photoreceptor cells remained intriguing and we decided to join forces to test the immunohistochemistry of CRY1 in the human retina, in which no convincing immunostaining has been reported so far. Our current research suggests that CRY1 is also located in the outer segments of blue cones (SWS1 cones) in hominid primates, including humans, bonobos, and gorillas and, furthermore, suggests a new explanation for why the antibodies used here specifically recognizes CRY1 in the SWS1 cones, and why CRY1 immunostainings can be challenging, inconclusive, and even contradictory.

## Results

We immunohistochemically detected the presence of CRY1 in the human retina as well as in the retinae of the Western lowland gorilla (*Gorilla gorilla gorilla*) and bonobo (*Pan paniscus*), using an antiserum that recognizes a sequence corresponding to the last 20 amino acids of CRY1 (gp-α-CRY1, Tables 1, 2). The CRY1 label (green) was apparent in the outer segments of a relatively sparse photoreceptor type (Fig. 1). Double labelling for the shortwave-sensitive (SWS1) cone opsin (magenta) showed that CRY1 is localized in the SWS1 (‘blue’) cones in human retina and also in the retinae of the other studied hominids. This includes the Sumatran orangutan (*Pongo abelii*), which we have studied before and show for comparison in Figure 1 ((31); there erroneously specified as Bornean orangutan *Pongo pygmaeus,* see correction (35)). The blue cones are present as a low-density population in human and great ape retinae (36, 37). The specific colocalization of CRY1 and SWS1 opsin labels in the blue cones is most clearly seen in the flatmount views in Fig. 1. The gp-α-CRY1 antiserum labelled no other retinal cell types, particularly no middle-to-longwave-sensitive (i.e. ‘green’ and ‘red’) cones containing the LWS opsins (Fig. 2). The human and bonobo retinae included both male and female samples, and the CRY1 label was the same in both sexes. The consistent CRY1 labelling across humans and great apes indicates that this is a common trait in the family Hominidae.

**Fig. 1.**
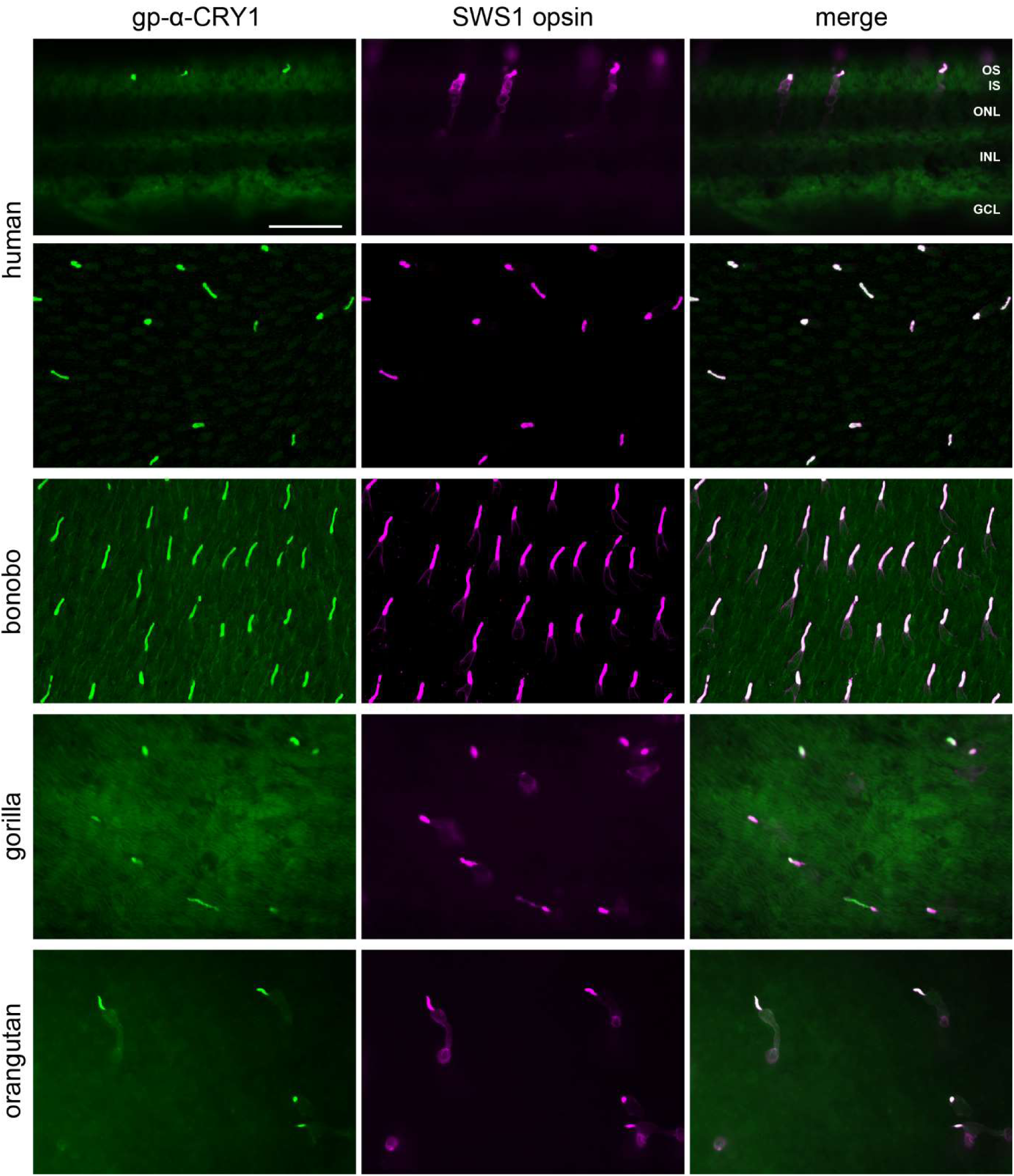
Full-length CRY1 label (detected with the gp-α-CRY1 serum) in the SWS1 (blue) cones of human, gorilla, bonobo, and orangutan retina. Transverse section (top) and flatmount of human retina, retinal flatmounts of the other species with focus on the photoreceptor layer. Left column: CRY1 immunofluorescence (green). Middle column: SWS1 opsin immunofluorescence located in the blue cone outer segments and to a lesser extent in the larger blue cone inner segments (magenta). Right column: Merged images, showing that CRY1 and SWS1 opsin co-localize in the blue cone outer segments. The bonobo field is from the central retina, the gorilla and orangutan fields are from the peripheral retina, and the human field is from an unknown location. OS, IS, photoreceptor outer and inner segments, respectively; ONL, outer nuclear layer; INL, inner nuclear layer; GCL, ganglion cell layer. Scale bar in the top left image is 50 µm and applies to all images.

**Fig. 2.**
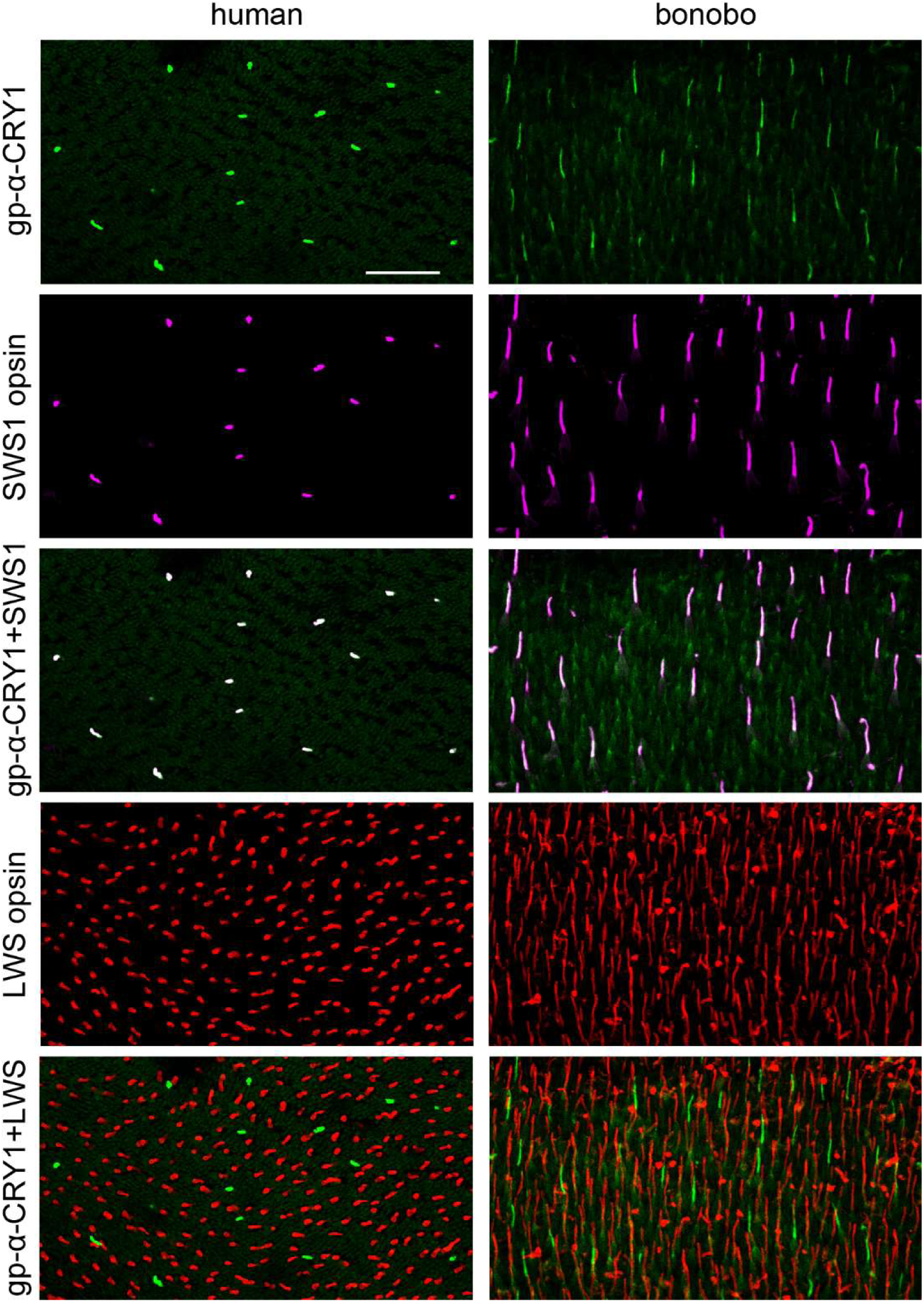
Triple-labelling for full-length CRY1 (gp-α-CRY1 serum), for the SWS1 opsin of the blue cones, and for the LWS opsin of the green and red cones in central human (left column) and bonobo retina (right column). Images from flatmounted retinal pieces, focused on the cone outer segments. In addition to the single-label images, merged images of the CRY1 label (in green) and the SWS1 opsin label (in magenta), and of the CRY1 label and the LWS opsin label (in red), are shown. The merged images confirm that the CRY1 label is limited to blue cones (whitish appearance) and does not occur in green and red cones (which would appear yellow). Photoreceptor outer segment preservation is less good in the human than in the bonobo tissue. The images are maximum intensity projections of confocal image stacks. Scale bar in the top left image is 50 µm and applies to all images.

**Table 1.** Primary CRY1 antisera / antibodies used in this study. Identical amino acids to human CRY1 are shown in red. IHC= Immunohistochemistry; WB = Western blot.

| antibody name | species and antibody clonality | immunization peptide | dilution | Source |
| --- | --- | --- | --- | --- |
| gp-α-CRY1 | guinea pig polyclonal | RPNPEEETQSVGPKVQRQST<br>hCRY1 sequence 566-585 | IHC:<br>1:100<br>WB:<br>1:500 | Genovac GmbH, Freiburg, Germany (32) |
| rb-α-CRY1 PA1-527 | rabbit polyclonal | QSVGPKVQRQSSN<br>hCRY1 sequence 574-586 | IHC:<br>1:500<br>WB:<br>1:1000 | Thermo Fisher Scientific,<br>Waltham, MA, USA |
| rt-α-CRY1 3E12 | rat monoclonal | RPNPEEETQSVGPKVQRQST<br>hCRY1 sequence 566-585 | IHC:<br>undiluted | Monoclonal Antibody Core Facility,<br>Helmholtz Zentrum München,<br>German Research Center for Environmental Health (GmbH),<br>Neuherberg, Germany<br>(Bolte et al. 2021) |
| rt-α-CRY1 17A2 | rat monoclonal | TTPVSDDHDEKYG<br>hCRY1 sequence 179-191 | IHC:<br>1:2<br>WB<br>1:1000 | This study;<br>Monoclonal Antibody Core Facility,<br>Helmholtz Zentrum München,<br>German Research Center for Environmental Health (GmbH),<br>Neuherberg, Germany |
| rb-α-CRY1 ARP59758_P050 | rabbit polyclonal | hCRY1 sequence 151-200 | IHC:<br>1:200 | aviva Systems Biology, San Diego, CA, USA |

**Table 2.** Amino acid sequence of the antigen used to generate gp-α-CRY1 and rt-α-CRY1 3E12 antibodies compared with the sequences of CRY1 in the studied Hominidae. Identical amino acids are shown in red. The numbers of the amino acids (AA) compared to the antigen sequence, as well as the full AA length of the respective CRY1 proteins are indicated below the species names.

| Species | FASTA Sequence | Sequence ID |
| --- | --- | --- |
| Epitope recognized by antiserum/antibody | RPNPEEETQSVGPKVQRQST |  |
| *Homo sapiens* |  |  |
| Human | RPSQEEDTQSIGPKVQRQST | NP_004066.1 |
| AA 566 – 585 (length: 586) |  |  |
| *Pan paniscus* |  |  |
| Bonobo | RPSQEEDTQSIGPKVQRQST | XP_003832588.1 |
| AA 566 – 585 (length: 586) |  |  |
| *Pongo abelii* |  |  |
| Sumatran orangutan | RASQEEDTQSIGPKVQRQST | XP_002823736.1 |
| AA 566 – 585 (length: 586) |  |  |
| *Gorilla gorilla gorilla* |  |  |
| Western lowland gorilla | RPSQEEDTQSIGPKVQRQST | XP_018894476.1 |
| Predicted AA 566-585 (length:586) |  |  |

CRY1 labelling of the hominid SWS1 cone outer segments was further confirmed with two additional C-terminal binding antibodies (bonobo and human shown in Figs. 3 and S1; gorilla and orangutan not illustrated). Of these, the monoclonal antibody (rt-α-CRY1 3E12, Fig. 3, also used in (33)) was directed against the same sequence as the gp-α-CRY1 but produced in a different species (rat). The polyclonal antibody is commercially available (rb-α-CRY1 PA1-527, Thermo Fisher Scientific) and recognizes a shorter sequence, but also the C-terminus (Fig. S1). These two antibodies also label the SWS1 cone outer segments, showing a pattern very similar to that of gp-α-CRY1. This strongly corroborates the statement that our immunostainings actually show CRY1 and not an off-target protein. The antibody rb-α-CRY1 PA1-527 labelled a small number of somata in other retinal layers, an example in the human inner plexiform layer is shown in Fig. S1. This may indicate that a few other retinal cells may contain full-length CRY1 with the epitope available to the antibody. However, it could also be a false positive signal, as Western blots with rb-α-CRY1 PA1-527 show a second unspecific band in HEK cells (Fig. S4).

**Fig. 3.**
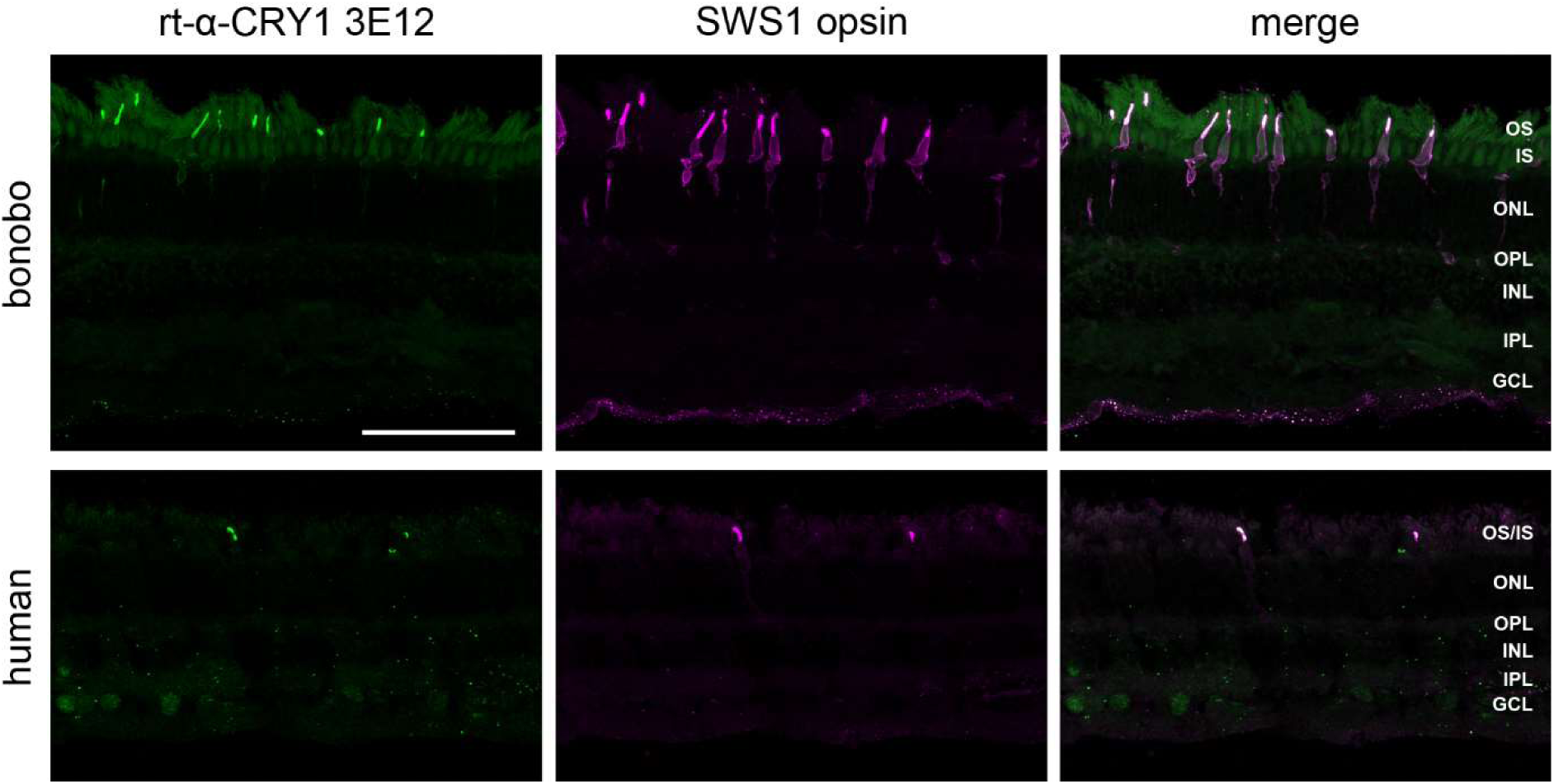
Full-length CRY1 label (detected with rt-α-CRY1 3E12) in the SWS1 (blue) cones of bonobo and human, retina in transverse sections. Left column: CRY1 immunofluorescence (green). Middle column: SWS1 opsin immunofluorescence located in the blue cone outer segments, and in bonobo also more faintly across the whole blue cones (magenta). Right column: Merged images, showing that CRY1 and SWS1 opsin co-localize in the blue cone outer segments. The human tissue preservation is less good. The images are maximum intensity projections of confocal image stacks. OS, IS, photoreceptor outer and inner segments; ONL, outer nuclear layer; OPL, outer plexiform layer; INL, inner nuclear layer; IPL, inner plexiform layer; GCL, ganglion cell layer. Scale bar in the top left image is 100 µm and applies to all images.

To further test the guinea pig serum’s specificity towards CRY1, we expressed human CRY1 (hCRY1) tagged with GFP in HEK cells. The Western blots in Figure 4 demonstrate that the antiserum was indeed able to recognize a band at the expected size of hCRY1-GFP (93 kDa) that was absent in the GFP only control. Smaller unspecific bands in the Western blots seemed to be the result of the secondary HRP goat anti-guinea pig antibody (Fig. 4B), which was not used in the retina immunohistochemistry. We furthermore blasted the antigen sequence to identify possible cross-reactivities. The only hit that made sense in the tissue context was the rod-specific cyclic nucleotide-gated cation channel beta 1 (CNGB1) subunit sharing eight amino acids with the antigen (Table 3). To test whether these eight amino acids were sufficient for recognition by the serum, we replaced the last 20 amino acids of CRY1 with 10 amino acids from CNGB1. Expression of this chimera (hCRY1^Δ566-585^-CNGB1^41-50^-GFP) in HEK cells confirmed the loss of recognition by the serum (Fig. 5). Notably, this CNGB1 sequence is not part of the antigen sequence recognized by rb-α-CRY1 PA1-527. These controls further confirm that the signal seen in the SWS1 cones is indeed CRY1. Two anti-CRY1 antibodies targeting N-terminal regions of CRY1 (rt-α-CRY1 17A2 produced in this study and the commercially available rb-α-CRY1 from Aviva) did not show selective labelling in the retina (Fig. S2). This could be because CRY2 may be the more relevant clock CRY in the retina, as suggested for birds (33, 38) and humans (18, 39). Furthermore, rt-α-CRY1 17A2 recognized the C-terminally truncated form of CRY1 (hCRY1^Δ566-585^-CNGB1^41-50^-GFP) much better than the full-length protein (Fig. 5). We suggest that the recognition sites of antibodies targeting the N-terminal region are partially blocked by the C-terminus of CRY1 binding to the PHR domain (12), and even the denaturing conditions of the SDS-PAGE seem to have not unfolded CRY1 completely, potentially explaining why the two N-terminal region recognizing antibodies did not stain full-length CRY1 in the SWS1 cones.

**Fig. 4.**
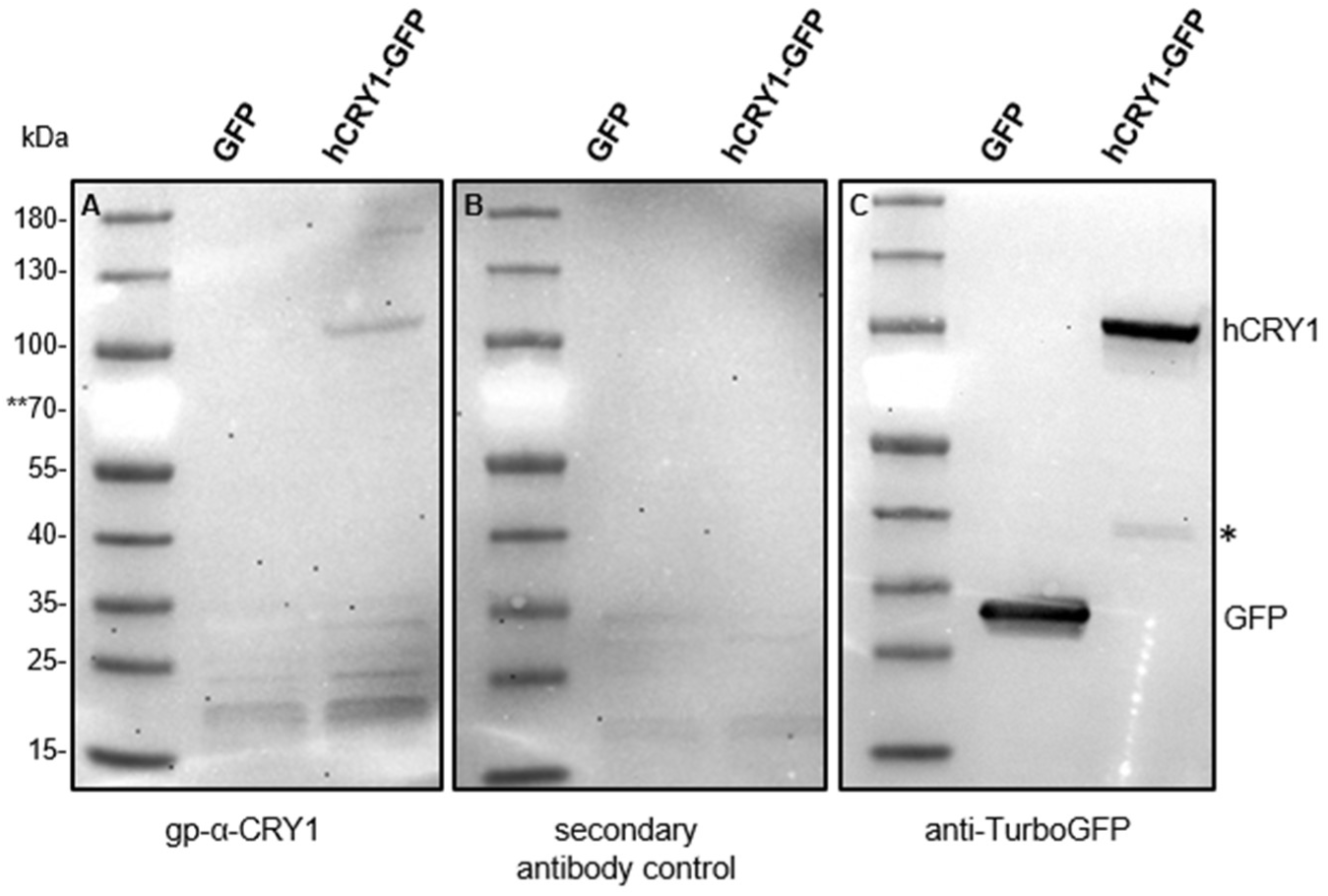
Western Blot of GFP and CRY1-GFP expressed in HEK 293 cells. 40 µg of HEK cell lysate expressing either GFP alone or a CRY1-GFP fusion protein was subjected to SDS-PAGE and Western blotting. A Western blot incubated with the gp-α-CRY1 serum shows a band close to the expected size of 93 kDa, but not in HEK cells expressing GFP only. Other unspecific bands seem to come from the secondary antibody, as demonstrated in the Western blot incubated with the secondary antibody only (B). A Western blot incubated with an anti-TurboGFP antibody was used as a positive control (C). * This band likely indicates a truncated C-terminus with the GFP-tag. ** The red colour of the 70 kDa protein marker appears bright white with illumination.

**Fig. 5.**
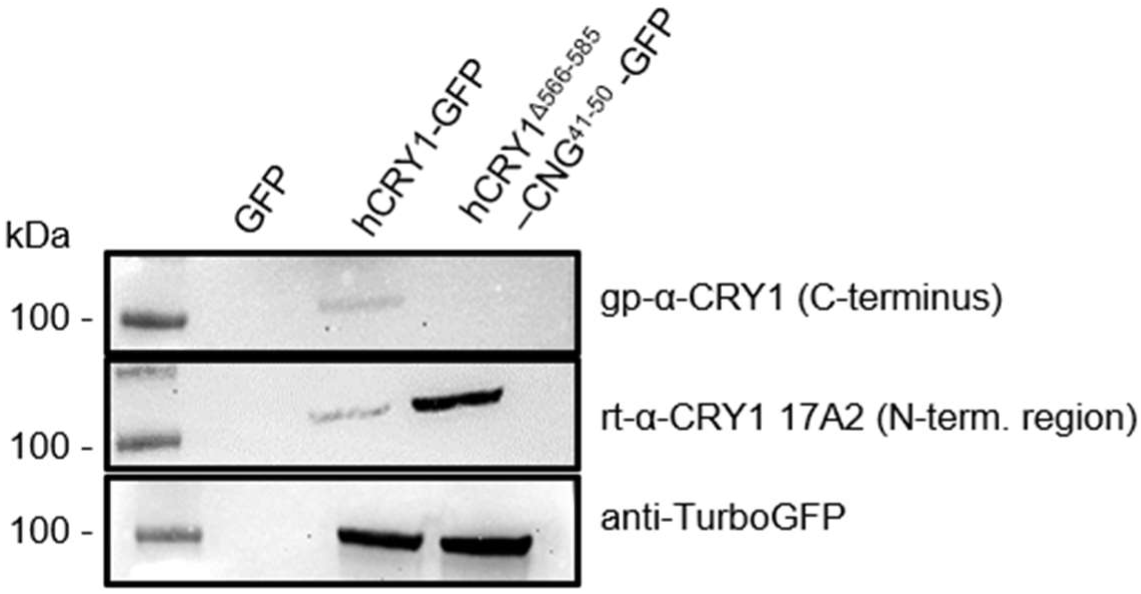
Western blot of hCRY1-GFP or hCRY1^Δ566-585^-CNGB1^41-50^-GFP expressed in HEK 293 cells. 40 µg of HEK 293 cell lysate expressing the empty plasmid (GFP), a CRY1-GFP fusion protein or a hCRY1^Δ566-585^-CNGB1^41-50^-GFP chimeric fusion protein were analysed using SDS-PAGE and Western blotting. The gp-α-CRY1 serum used in this study recognized only full-length CRY1, but not the hCRY1^Δ566-585^-CNGB1^41-50^-GFP chimeric protein. Interestingly, the monoclonal rt-α-CRY1 17A2 directed against an N-terminal region of CRY1 recognized the truncated CRY1 protein much better than the full-length protein. A control staining against TurboGFP indicates that both proteins were expressed in approximately equal amounts. Full Western bot images can be found in the supplements (Fig. S3).

**Table 3.** Amino acid sequence of the antigen recognized by gp-α-CRY1 compared to the sequence of the rod-specific CNG-channel beta 1 subunit. Identical amino acids are shown in red.

| Species | FASTA Sequence | Sequence ID |
| --- | --- | --- |
| Epitope recognized by gp-α-CRY1 | RPNPEEETQSVGPKVQRQST | |
| *Homo sapiens* CNG-channel beta and gamma subunit | PNPEEAETES | AAC04830.1 |

As CRY1 is primarily known as a transcriptional repressor (4, 6–8), we wondered why we did not see any nuclear stainings in the retina and performed subcellular fractionations of hCRY1 containing a small N-terminal His-tag expressed in HEK cells. As the gp-α-CRY1 serum was limited, we analyzed these Western blots with the commercially available C-terminal antiserum rb-α-CRY1 PA1-527 (Fisher Scientific) (Table 4). Subcellular fractionations demonstrated a signal for full-length hCRY1 in the cytosol and membrane fraction, but not in the nuclear fraction (Fig. 6). An immunofluorescence analysis antibody test by Fisher Scientific for PA1-527 in U-87 MG cells also shows cytosolic, but no nuclear labelling (www.thermofisher.com/antibody/product/CRY1-Antibody_polyclonal/PA1-527). Furthermore, a literature search failed to find any nuclear labelling of CRY1 with a defined C-terminal antibody.

**Fig. 6.**
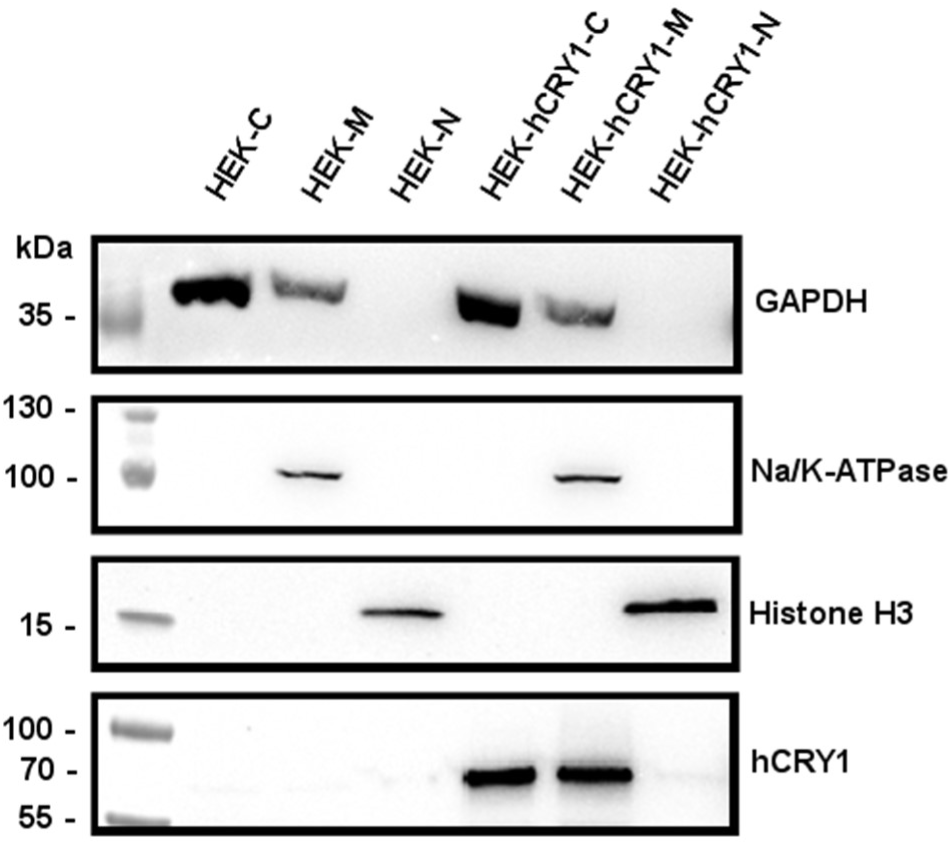
Subcellular fractionation of HEK cells with and without transfected hCRY1. Cell lysates of the cytosolic fraction (C), membrane fraction (M) and nuclear fraction (N) were subjected to SDS-PAGE and Western blotting and probed against GAPDH as a marker for cytosolic and membrane fractions, against Na/K-ATPase as a marker for the membrane fraction and against Histone H3 as a nuclear marker. The CRY1 antiserum used in this Western blot was rb-α-CRY1 PA1-527, recognizing full-length CRY1. Full Western bot images can be found in the supplements (Fig. S4).

**Table 4.** Amino acid sequence of the antigen recognized by rb-α-CRY1 PA1-527. Identical amino acids are shown in red. The numbers of the amino acids (AA) compared to the antigen sequence, as well as the full AA length of the human CRY1 protein are indicated below the species names.

| Species | FASTA Sequence | Sequence ID |
| --- | --- | --- |
| Epitope recognized by rb-α-CRY1 PA1-527 | QSVGPKVQRQSSN | |
| *Homo sapiens* |  |  |
| Human<br>AA 574 – 586 (length: 586) | QSIGPKVQRQSTN | NP_004066.1 |
| *Pan paniscus* |  |  |
| Bonobo<br>AA 566 – 585 (length: 586) | QSIGPKVQRQST | XP_003832588.1 |

## Discussion

### Localization in the outer segments of SWS1 cones suggests an additional functional role for full-length CRY1 in the retina

Light influences our health and quality of life in many ways, including entraining our circadian rhythm, controlling the pupillary light reflex, and affecting sleep and mood. In mammals, the only entrance point for light to image-forming and non-image-forming visual processes is the retina (reviewed in (40)). Melanopsin containing retinal ganglion cells were found to play a significant, but not exclusive part in mediating non-image-forming photoresponses (21, 41). Besides melanopsin, CRYs have also been suggested to be involved (15, 27, 42, 43). However, the localization of CRYs in the human retina remains a matter of debate, especially because CRY1 had not been detected on a protein level until now. Here, three different C-terminal directed CRY1 antibodies suggest a rather specific localization of full-length CRY1 in the SWS1 (blue) cone photoreceptor outer segments, uncovering a previously overlooked aspect of CRY1’s distribution and function within the human and great ape retinae. While past investigations have primarily focused on the involvement of CRY1 in circadian rhythm regulation, particularly within the nucleus, it is unlikely that full-length CRY1 solely functions as a transcriptional repressor in these cells, given the compartmentalization of photoreceptors with multiple barriers between the outer segment and the nucleus (44). In particular, the absence of any nuclear staining raises the intriguing possibility that full-length CRY1 could be specific to SWS1 cones (the blue cones of mammals, and CRY1a to the UV/violet cones of birds) in the retina, potentially serving a unique functional role. It is interesting to note that the outer segments could be stained regardless of the time of death of the donors (which is unknown), which could mean that the expression might be independent of the circadian rhythm in these cells, or a lack of degradation.

Remarkably, the mouse retina does not show an equivalent immunohistochemistry of CRY1. Both the same gp-α-CRY1 serum as well as the rb-α-CRY1 PA1-527 antibody used in this study did not label any cone photoreceptors in the mouse in two independent studies (21, 31). We therefore may need to reconsider whether the predominantly nocturnal mouse is an appropriate model species for CRY1 in the human retina.

In contrast to the nuclei, the outer segments of photoreceptors are specialized for photoreception and contain all the necessary cellular machinery for phototransduction. If CRY1 binds its FAD cofactor in these cells (but see (24)), CRY1 would most effectively absorb light in the blue part of the spectrum, aligning with the wavelength sensitivity of the blue opsins. One could therefore speculate that CRY1 could regulate or enhance light detection in the SWS1 cones. Another possibility would be that CRY1 has a function entirely separated from the blue opsin, but that such a separate function requires blue light as well.

Discussions of CRYs as potential components of the light-dependent magnetic compass in birds (25, 45–49) underscore CRY’s diverse functionalities. However, of all avian CRYs, only CRY4 has thus far been successfully purified with FAD bound and demonstrated to be magnetically sensitive *in vitro* (25). Evidence for any involvement of CRY1 or CRY2 in magnetoreception is controversial (24, 25, 33, 50–52). Critically, the suggestion that hCRY2 can rescue light-dependent magnetosensation in *Dm*Cry-deficient *Drosophila* (53) has recently been seriously challenged in an independent replication attempt that found no evidence for an influence of magnetic fields on the behaviour of fruit flies in the first place (54, 55). Thus, mammalian CRY’s role in magnetoreception remains uncertain, but the localization of hCRY1 in the outer segments could allow for an involvement in processes related to phototransduction. Interestingly, an interaction between avian CRY1a/b and the cone-specific G-protein (transducin) alpha subunit, which is involved in the phototransduction cascade, was observed, albeit one order of magnitude weaker than the strong interaction between the light-sensitive putative magnetoreceptor CRY4 and cone transducin (56). If CRY1 in SWS1 cones interacts similarly, this could indicate a role in initiating or modulating light-induced signal transduction, or a means of anchoring cryptochromes or mediating their trafficking to the outer segments (57).

Alternatively, hCRY1 in SWS1 cone photoreceptors may have a possible role in sensing or dealing with oxidative stress. In contrast to medium- and long-wavelength sensitive cones, SWS1 cones appear to have limited capacity for aerobic glycolysis and rely largely on oxidative phosphorylation (58), which is known to generate reactive oxygen species. hCRY1 in transgenic *Drosophila* has been linked to the regulation of genes implicated in stress response and reactive oxygen species (ROS) signaling (59), but the precise mechanisms remain to be clarified. In summary, the localization of full-length CRY1 in SWS1 cones challenges traditional views of the exclusive role of CRY1 as a nuclear protein and supports additional functions which could include or affect phototransduction or light-sensing mechanisms within the retina, a function that may be unique to the full-length version of CRY1.

### CRY1 immunostainings can be misleading

Binding of two C-terminal specific antibodies to full-length CRY1 was only detected in the outer segments of SWS1 cones with no corresponding labelling detected in other retinal cell types. It is, nevertheless, still possible that full-length CRY1 exists elsewhere, but that the binding site for the antibodies is hidden by interaction partners or by different folding of the C-terminus, which has been notoriously difficult to crystallize in most cryptochromes (60–63). A third antibody, rb-α-CRY1 PA1-527, labelled, in addition to the SWS1 cones, also a few cells in other retinal layers, but the specificity of this antibody is questioned by Western blots. The outer segment localization of CRY1 seems surprising, not only in the light of its known clock function, but also when considering that the *CRY1* gene was found to be expressed across the entire retina, albeit at significantly lower mRNA levels than *CRY*2 (18, 39). However, Thompson et al. (18) reported no detection of CRY1 protein. We also did not detect selective labelling across the retinal layers using two different antibodies against the N-terminal region of CRY1 (Fig. S2), which either means that these antibodies are unspecific or a C-terminal truncated version of CRY1 is found in many retinal cells. Importantly, our CRY1 antibodies that specifically label SWS1 cones target the C-terminal sequence of full-length CRY1 and do not recognize shorter versions with truncated C-terminal sequences, whose presence we cannot exclude in the retina. We did not find out which CRY1 antibodies were unsuccessfully tested by Thompson et al. (18), but in our hands only C-terminally directed antisera showed a specific immunoreactivity in the human retina.

Our Western blot analysis indicates that the rt-α-CRY1 17A2 directed against the N-terminal region of CRY1 preferably recognizes a truncated version of CRY1, suggesting it is unable to recognize the native full-length CRY1 protein in immunohistochemistry. The reason may be that the unknown way in which the C-terminus binds to the PHR domain (12), might block the binding site. If other N-terminal region antibody binding sites are also blocked by the C-terminus, the consequences are that immunostainings performed with only one antibody against CRY1 will not show the complete picture, thus explaining exclusive and non-overlapping stainings of N- and C-terminal recognizing CRY1 antibodies, questioning many conclusions based on CRY1 immunostainings alone. Thus, CRY1 antibodies need to be carefully validated, which is not trivial, as small amino-acid losses in CRY1’s C-terminal region may go unnoticed even in Western blots, but may have big consequences for antibody recognition, especially in immunohistochemistry. We can therefore only conclude that full-length CRY1 can, so far, only be documented in the SWS1 cones in the hominid retina. However, we cannot exclude that other (shorter) CRY1 versions exist in a perhaps oscillating manner in other retinal cell types, or that full-length CRY1 exists elsewhere with its binding site not accessible to the antibody.

### The role of CRY1’s C-terminus

CRY1 is often described as “instable”, depicted in Western blots as a 55 kDa version (64), or as a double band (13, 65) instead of a single band at its expected size of at least 66 kDa. Also, our own Western blot experiments on CRY1-GFP (Fig. 4c) show a second specific band recognized by the GFP antibody, which is much smaller (∼37 kDa) than the full-length protein with GFP (93 kDa), but larger than GFP alone (27 kDa). The most obvious suggestion is that the band at ∼37 kDa could be a truncated CRY1 C-terminus-part with GFP attached. The observation of truncated CRY1 protein in Western blots along with the unexpected localization pattern of full-length CRY1 in photoreceptor outer segments in retinal tissues suggests potentially different roles of CRY1, maybe based on different, more or less truncated versions of CRY1. Parts of the C-terminal amino acid sequence could be targeting CRY1 to the outer segments of photoreceptor cells, similar to the C-terminal tail of rhodopsin, where a rod outer segment localization signal resides within the terminal eight amino acids (66). Intriguingly, avian cryptochrome 1a and 1b, which differ only in their C-terminus with CRY1b having a very different and markedly shorter C-terminus, show a different localization in the avian retina: While the same gp-α-CRY1 serum as used here localized avian CRY1a to the UV/V cone outer segments (32, 33), CRY1b was found in ganglion cells, displaced ganglion cells, and photoreceptor inner segments of the avian retina (67).

This prompts the question, whether the protein thought to play a central role in the circadian clock is the full-length CRY1, a shorter version, or both. While the answer to this is beyond the scope of this paper, several observations argue (a) that CRY1 more easily forms a complex with PER without its C-terminus (12), and (b) that the C-terminal tail of CRY1 seems to be dispensable for its repressive function on CLOCK:BMAL1 (68, 69). Intriguingly, a 55 kDa version of CRY1 accumulates in the nuclear fraction of outer root sheath cells of hair follicles upon light exposure (64), suggesting the protein has “lost” approximately 11 kDa. In our study, we neither observed any full-length CRY1 in the nuclei of the retina, nor in the nuclear fraction of HEK cells, indicating further that the known nuclear circadian clock protein is perhaps a shorter version of CRY1.

Several studies show that ubiquitination and degradation of CRY1 regulate the molecular clock (70–72), but whether a proteolytic processing of CRY1’s C-terminus is specifically needed in this mechanism is unknown. We do know, however, that phosphorylation of CRY1’s C-terminus modulates circadian period length (13). Our study stresses the importance of the exact length of CRY1 and the consequences this has for antibody recognition and the circadian clock mechanism.

These discoveries open avenues for further research into the functional significance of CRY1’s C-terminus, and its putative role in visual perception, or another new, entirely unsuspected function. Such investigations are likely to broaden our understanding of CRY1’s role in circadian regulation and beyond.

## Materials and Methods

### Tissue sources

Retinal tissue of 12 human eyes from 12 donors (six males, six females; age range 42-89 years) was used for this study. Some retinae were provided after enucleation of the eye by university hospitals in Tübingen (Ethics Commission of the Tübingen University Medical Faculty, approved by permit number 531/2011BO2). Other retinae were provided *post mortem* by the Cornea Bank of Rhineland-Palatine, Department of Ophthalmology, University Medical Center Mainz (organ donations followed the guidelines for corneal donations regulated by the German Transplantation Law; the use of remaining ocular donor tissue not required for transplantation for study purposes has been explicitly approved by the relatives of the deceased and the local ethics committee), and by the Amsterdam Cornea Bank via the Institut de la Vision, Paris, France (French Ministère de l’Education et de la Recherche Scientifique, approved by permit number DC-2015-2400). All human tissue was obtained with the informed consent of the donors or their relatives, respectively. No tissues were procured from prisoners. All methods were performed in accordance with all relevant guidelines and regulations.

The eyes of a 32 years old male and a six months old female bonobo (*Pan paniscus*), a 51 years old female Western lowland gorilla (*Gorilla gorilla gorilla*), and a 57 years old male Sumatran orangutan (*Pongo pygmaeus abelii*) were used; they were obtained at autopsies when the animals had died in Frankfurt Zoo. Tissue use for research was permitted by CITES certificates DE-DA-240327-1 for the adult male bonobo, DE-DA-240327-2 for the juvenile female bonobo, DE-DA-160824-7 for the gorilla, and DE-DA-240708-9 (Senckenberg Research Institute Section Mammalogy collection number SMF 98848) for the orangutan. The orangutan tissue had already been used in our previous study (31). None of the human and great ape eyes had known retinal pathologies.

### Tissue preparation

The human eyes were opened by a cut around the cornea, and the retina was isolated and immersion-fixed in 4 % paraformaldehyde in 0.1 M phosphate buffer (PB, pH 7.4) or 0.01 M phosphate-buffered saline (PBS, pH 7.4). Fixation time of the human retinae was between 20-30 min and 12 h. After fixation, the tissue was stored in PBS at 4°C until further use. The gorilla and bonobo eyes were obtained several hours *post mortem*, opened by a cut around the cornea and immersion-fixed in 4% paraformaldehyde in PB for one day (gorilla eyes) or one month (bonobo eyes). After washing out the fixative with several changes of PB, the retinae were isolated. Whole retinae or retinal pieces were cryoprotected in an ascending series of 10%, 20% and 30% sucrose in PB and frozen at -20°C for storage. We used both 16 µm thick cryosections and pieces of unsectioned retinae (flatmounts) for staining.

Enucleated human eyes were exposed to bright light in the operating theatre, but then kept in darkness for transport. Retinae were isolated in red light and fixed in darkness, after fixation they were exposed to various light conditions. To our knowledge, the bonobo, gorilla and orangutan eyes had been exposed to white light (daylight or laboratory lighting) before and during fixation.

### CRY1 Antibodies

The gp-α-CRY1 antiserum used previously (31, 32, 73), was no longer available and a new lot of the antiserum was ordered and validated here in cell culture, see details further below. A new monoclonal antibody production for rt-α-CRY1 17A2 was performed as described in (47). The peptide for immunization was produced by Peps 4LS GmbH (Heidelberg, Germany). Details on all CRY1 antibodies used in this study are listed in Table 1.

### Immunohistochemistry

The retinae were pre-incubated with 10% normal donkey serum (NDS) in 0.5% Triton X-100, 1% bovine serum albumin (BSA) in PB for 60 min at room temperature (RT). Then they were incubated in a mixture of one of the CRY1 antibodies (see Table 1) and the goat SWS1 opsin antiserum sc-14363 (1:500; Santa Cruz Biotechnology Inc., Santa Cruz, CA, USA) in 3% NDS, 0.5% Triton X-100, 1% BSA, in PB overnight for sections and 2-3 days for unsectioned retinal pieces. After washing in PB, the tissue was incubated in a mixture of the secondary antibodies donkey-anti-guinea pig IgG coupled to Alexa488 and donkey-anti-goat IgG coupled to Alexa649 (dilutions 1:500; Dianova, Hamburg, Germany) in 3% NDS, 0.5% Triton X-100, 1% BSA, in PB for 60 – 90 min.

For triple immuno-fluorescence labelling of CRY1, SWS1 opsin, and LWS (‘red’ and ‘green’ cone) opsin, retinal pieces were incubated in a mixture of gp-α-CRY1, sc14363 and the rabbit LWS opsin antiserum JH492 (1:2000; kindly provided by Jeremy Nathans). Labelling was visualized with a mixture of the above-mentioned secondary antibodies and donkey-anti-rabbit IgG coupled to Cy3 (dilution 1:250; Dianova, Hamburg, Germany). The two cone opsin antisera sc-14363 and JH492 have been shown to robustly label the SWS1 opsin and LWS opsin, respectively, across mammals (see (31) and references therein).

After staining, the retinae were coverslipped with an aqueous mounting medium and evaluated at a Zeiss Axioplan 2 microscope using the Axiovision LE software (Carl Zeiss Vision), and at a laser scanning microscope (LSM) Olympus FluoView 1000 using the FV 1.7 software (Olympus). LSM images and z-stack projections were examined with ImageJ (https://imagej.net). Images for illustration were adjusted for brightness and contrast using Adobe Photoshop.

### Cloning

Human *CRY1* cDNA was amplified from pDONR223_hCRY1_WT, a gift from Jesse Boehm & William Hahn & David Root (74) (Addgene plasmid # 82264; http://n2t.net/addgene:82264 ; RRID:Addgene_82264) using CloneAmp HiFi PCR Premix (Takara Bio Inc. Shiga, Japan). For the cloning of pTurbo-hCRY1-GFP, the sense primer 5’-GGACTCAGATCTCGAGCCACCATGGGGGTGAACGCCGT-3’and antisense primer 5’-GGCGACCGGTGGATCCCCATTAGTGCTCTGTCTCTGGACTTTAGG-3’were used to amplify h*CRY1*. The purified PCR product was used for an In-Fusion reaction (Takara Bio Inc. Shiga, Japan) with *Xho*I and *BamH*I linearized pTurbo vector (Evrogen, Moscow, Russia). To generate pTurbo-hCRY1^Δ566-585^-CNGB1^41-50^-GFP, the C-terminal sequence of h*CRY1* corresponding to the antiserum recognition site (amino acids 566-585) was replaced with a ten amino acid sequence from CNGB1 (PNPEEAETES) using the Q5 site-directed mutagenesis kit (New England Biolabs, Ipswich, MA, USA).

For the cloning of pcDNA3.1-His-hCRY1, h*CRY1* was amplified using sense primer 5’-ATATGCTAGCGGCCATTACGGCCATGGGGGTGAACGCCG-3’ and antisense primer 5’-GCAGAATTCTGGCCGAGGCGGCCCTAATTAGTGCTCTGTCTCTGGACTTTAGG-3’.

The purified PCR product was then cloned into the *Sfi*I linearized vector pcDNA3.1^(+)^ (Thermo Fisher Scientific, Waltham, MA, USA) modified to contain a Kozak sequence, an N-terminal deca-histidine tag and an *Sfi*I restriction site (a gift from Prof. Karl Koch, Carl von Ossietzky University Oldenburg), using In-Fusion Master Mix (Takara Bio Inc. Shiga, Japan). All sequences were confirmed via Sanger sequencing (LGC Genomics).

### Cell Culture and Transfections

Human embryonic kidney 293 (HEK293) cells (ECACC 85120602) were cultured in DMEM + GlutaMax (Gibco, Waltham, MA, USA) supplemented with 10% fetal bovine serum (Gibco) at 37°C and 5% CO_2_. 2 × 10^6^ HEK cells were seeded in a 6-cm petri dish and transfected the next day using 7.5 µg of plasmid and 5 µl of Lipofectamine 2000 (Thermo Fisher Scientific, Waltham, MA, USA) per dish according to the manufacturer’s instructions. The medium was exchanged 24 h after transfection and cells were collected 48 h post transfection.

### Subcellular fractionation

Subcellular fractionation of transfected HEK cells was conducted following the protocol provided by Baghirova et al. (75): 48 h post transfection, the culture medium was removed and cells were washed with a phosphate buffered saline solution (PBS) (Gibco, Waltham, MA, USA). Cells were then trypsinized using 500 µl of 0.5% Trypsin-EDTA (Gibco, Waltham, MA, USA) for 1 minute at 37°C. 2 ml of culture media were added to inhibit trypsin activity and collect the cells. Cells were counted using Countess II FL (Thermo Fisher, Waltham, MA, USA), centrifuged for 10 min at 500 x g and 4°C, washed in ice-cold PBS and centrifuged again at 1000 x g for 20 min at 4°C, shock-frozen in N_2_ and stored at -80°C for up to one week. 5 × 10^6^ cells were lysed in 400 µl ice-cold lysis buffer A (50 mM HEPES, pH 7.4, 150 mM NaCl, 25 µg/ml digitonin and 1 M hexylene glycol) supplemented with protease inhibitor cocktail (cOmplete Mini EDTA-free tablets, Roche, Basel, Switzerland) for 10 min at 7°C rotating, and collected for 10 min at 4°C and 2000 x g. The supernatant contained the cytosolic fraction. The pellet was resuspended in 400 µl lysis buffer B (50 mM HEPES, pH 7.4, 150 mM NaCl, 1% Igepal, 1 M hexylene glycol) supplemented with protease inhibitor cocktail for 30 min rotating at 7°C and collected for 10 min at 4°C and 7000 x g. The supernatant contained membrane-bound organelles (mitochondria, endoplasmic reticulum, Golgi, etc.). The pellet was resuspended in 400 µl ice-cold lysis buffer C (50 mM HEPES, pH 7.4, 150 mM NaCl, 0.5% sodium deoxycholate, 0.1% sodium dodecyl sulfate, 1 M hexylene glycol) supplemented with 5 µl Benzonase and protease inhibitor cocktail, incubated for 40 min rotating at 7°C and collected for 10 min at 4°C and 7800 x g. The supernatant of this fraction contained nuclear proteins.

### Western blots

Cell pellets that were not used for subcellular fractionations were resuspended in 50 µl 20 mM Tris, pH 7.4, 150 mM NaCl and protease inhibitors and lysed by four cycles of freeze-thawing in liquid N_2_, and centrifuged for 10 min at 6000 x g and 4°C. All supernatant fractions were mixed with SDS sample buffer to a final concentration of 50 mM Tris, pH 6.8, 2.5% SDS, 0.02% bromophenol blue, 10% glycerol, and 1% β-mercaptoethanol and incubated for 5 min at 95°C (cytosolic fraction) or for 30 min at RT (membrane and nuclear fractions). SDS-PAGE was performed either with standard gels (10 – 12% acrylamide depending on protein size with a 4% acrylamide stacking gel) or 4-15% Mini-Protean TGX precast protein gels (Bio-Rad, Hercules, CA, USA) for CRY1 and Histone at constant 160 V. Proteins were transferred to an Amersham Protran Premium 0.2 NC nitrocellulose Western blotting membrane (Cytiva, Marlborough, MA, USA) using a Trans-Blot® Turbo^TM^ Transfer System (Bio-Rad Laboratories, Hercules, USA) and a gradient transfer buffer (0.3 M Tris, 20% methanol anode buffer and 24 mM Tris and 32 mM ε-amino n-caproic acid cathode buffer) for 30 min at 25 V. After blocking the membranes for one hour in 5% milk powder in TBST (20 mM Tris pH 7.4, 150 mM NaCl, 0.1% Tween 20), they were incubated with the primary antibodies (for details see Table 1 and Table S1) in 2.5% milk powder in TBST over night at 7°C rotating. Membranes were washed three times for 10 minutes in TBST and incubated with the respective secondary antibody (listed in Table S1) for one hour at RT. After again three washes in TBST for 10 minutes, membranes were developed using Super Signal West Pico chemiluminescent substrates (Thermo Fisher Scientific, Waltham, MA, USA) and visualized with the iBright CL 1000 (Thermo Fisher Scientific, Waltham, MA, USA).

## Acknowledgments

We are grateful to Dr. Deniz Dalkara (Institut de la Vision, Paris), Dr. Norbert Pfeifer (University Clinic, JGU Mainz), and Prof. Karl Ulrich Bartz-Schmidt and the surgery team of the University Clinics Tübingen for making the human retinae available to us, and to Dr. Christina Geiger and Dr. Nicole Schauerte (Zoo Frankfurt) and Dr. Kerstin Mätz-Rensing (German Primate Center Göttingen) for making the gorilla and bonobo eyes available to us. We are grateful to Prof. Karl W. Koch (Carl von Ossietzky University Oldenburg) for providing a plasmid and valuable discussions. Dr. Jeremy Nathans (Johns Hopkins Medical School Baltimore) kindly provided the LWS opsin antiserum JH492. We acknowledge the technical assistance of G. Heiß-Herzberger and G. Stern-Schneider in cryo-sectioning of retinae.

## Author contributions

R.B., C.N., L.P., and M.W. designed the study. R.B. performed the in-vitro work. C.N. and L.P. performed the histological stainings. P.B., R.F., K.D. and H.M. produced or prescreened CRY1 antisera or antibodies. R.B., C.N., L.P., and S.M. analyzed the data. K.R., H.M., and U.W. provided resources. R.B., C.N., L.P., and M.W. wrote the manuscript, with detailed feedback from K.D. and H.M. All authors discussed the results and contributed to the manuscript.

## Data availability

All relevant data are available in the paper.

