## Supplements for "Full-length Cryptochrome 1 in the outer segments of the retinal blue cone photoreceptors in humans and great apes suggests a role beyond transcriptional repression"

Rabea Bartölke[1*] (ORCID-ID 0000-0003-2531-747X), Christine Nießner (ORCID-ID 0000-0001-7985-4102)[2,3], Katja Reinhard (ORCID-ID 0000-0002-8719-7445)[4,5,#a], Uwe Wolfrum (ORCID-ID 0000-0002-4756-5872)[6], Sonja Meimann[7], Petra Bolte[1] (ORCID-ID 0000-0002-5514-6205), Regina Feederle (ORCID-ID 0000-0002-3981-367X)[8], Henrik Mouritsen (ORCID-ID 0000-0001-7082-4839)[1,10], Karin Dedek (ORCID-ID 0000-0003-1490-0141)[1,10], Leo Peichl (ORCID-ID 0000-0002-8141-0911)[2,3,7,9*], Michael Winklhofer (ORCID-ID 0000-0003-1352-9723)[1,10*]

*To whom correspondence should be addressed:

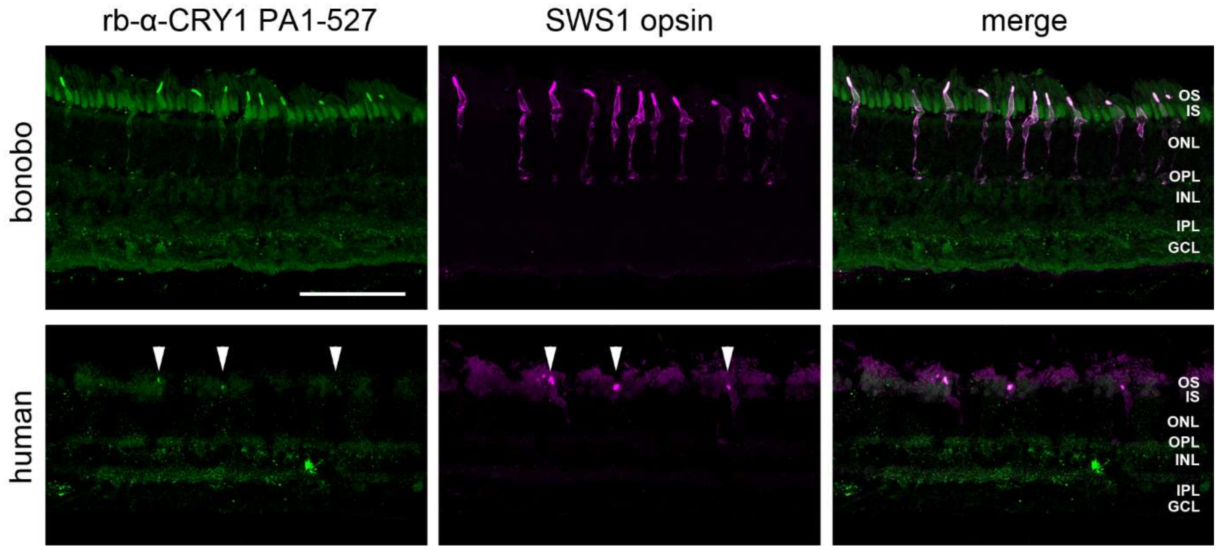


**Fig. S1** Full-length CRY1 label (detected with rb-α-CRY1 PA1-527, Thermo Fisher Scientific) in the SWS1 (blue) cones of bonobo and human retina in transverse sections. Left column: CRY1 immunofluorescence (green). The fainter green fluorescence in all cone inner segments and other retinal layers is unspecific background label by the secondary antibody; the strong label of an INL soma in the human section may or may not be specific. Middle column: SWS1 opsin immunofluorescence located in the blue cone outer segments, and in bonobo also across the whole blue cones (magenta). Right column: Merged images, showing that CRY1 and SWS1 opsin co-localize in the blue cone outer segments. The human tissue preservation is less good, so the blue cone outer segments (marked by arrow heads) are not as elongated as expected. The images are maximum intensity projections of confocal image stacks. OS, IS, photoreceptor outer and inner segments; ONL, outer nuclear layer; OPL, outer plexiform layer; INL, inner nuclear layer; IPL, inner plexiform layer; GCL, ganglion cell layer. The scale bar in the top left image is 100 µm and applies to all images.

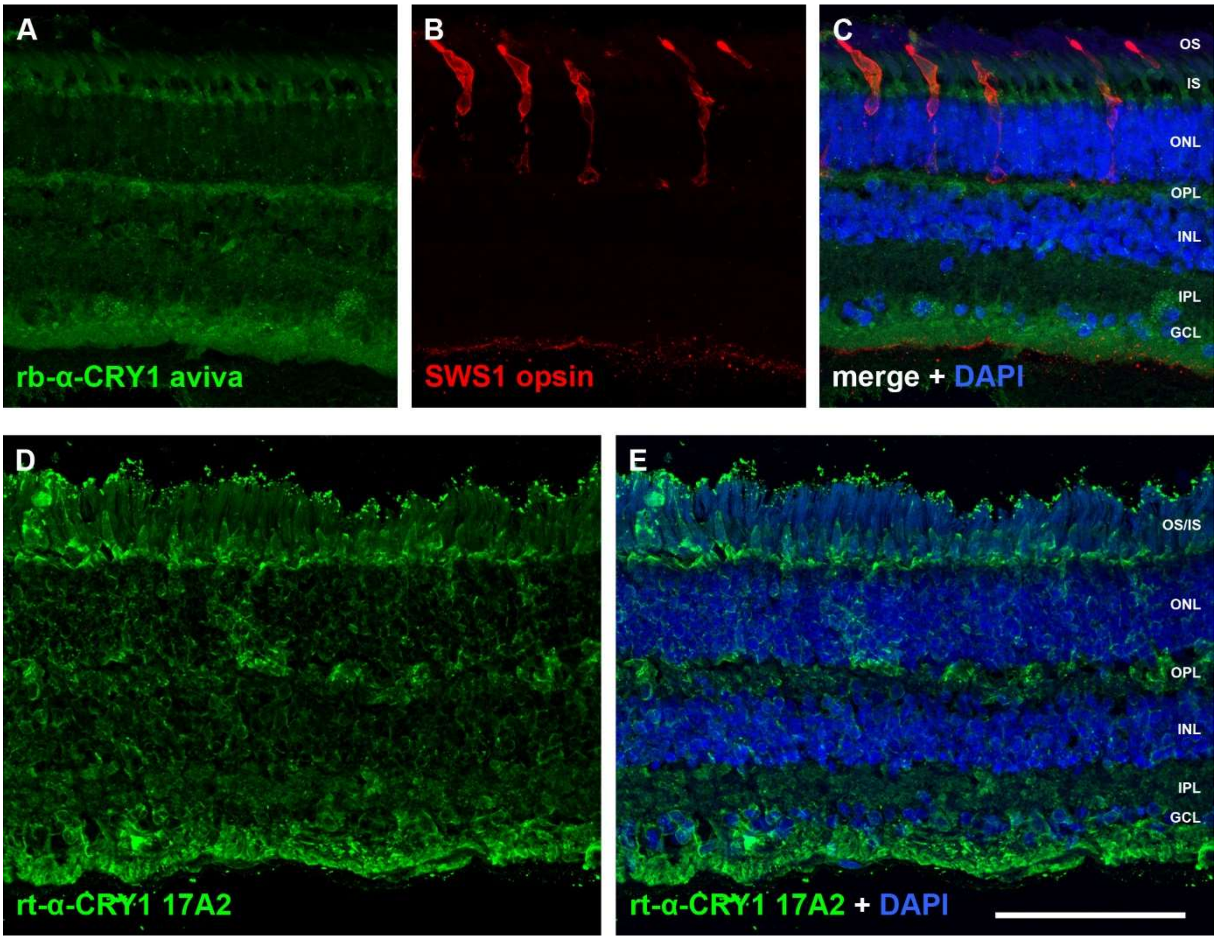


**Fig. S2** Absence of selective CRY1 label in SWS1 cones with antibodies directed against the N-terminus of CRY1. The images show transverse sections of bonobo retina as examples. (A-C) Double-labelling with the aviva rb-α-CRY1 antiserum ARP59758_P050 (A) and the SWS1 opsin antiserum (B); (C) merge of the two labels with DAPI counterstaining of the nuclear layers. This N-terminal-directed CRY1 antiserum shows no SWS1 cone outer segment label. (D, E) The N-terminal-directed rt-α-CRY1 antibody 17A2 also shows unselective labelling throughout the retina, but no specific cone labelling (D); (E) DAPI counterstaining. The images are maximum intensity projections of confocal image stacks. Retinal layers indicated as in Fig. S1. The scale bar is 100 μm and applies to all images.

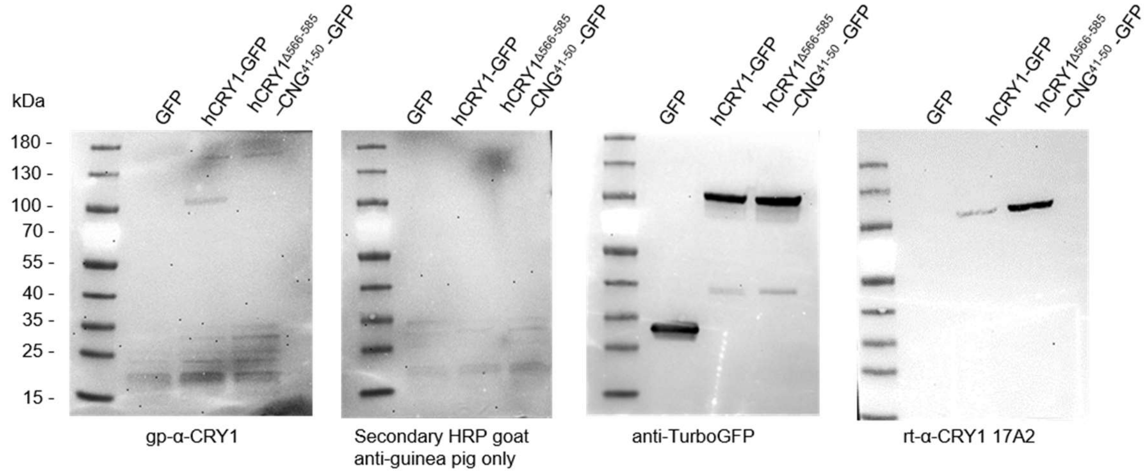


**Fig. S3** Full Western blot images of Figs. 4 and 5

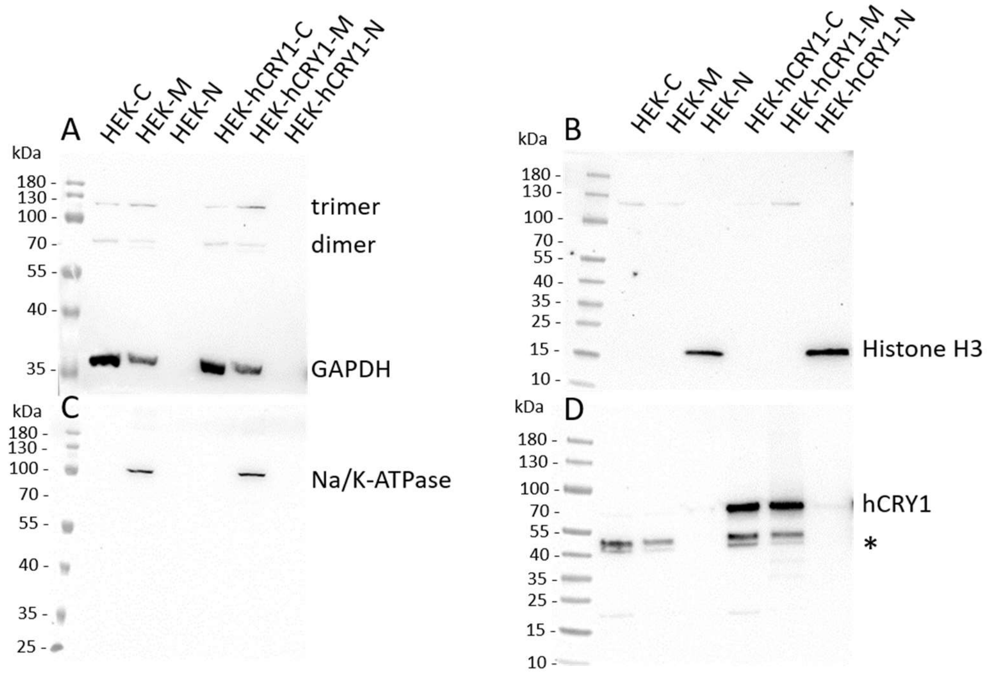


**Fig. S4** Full Western blot images of Fig. 6; Subcellular fractionation of HEK cells with and without transfected hCRY1. * unspecific band

**Table S1** Further antibodies used for Western blotting

| **antibody** | **species** | **dilution** | **Company** |
|---|---|---|---|
| anti-histone H3 (H0164) | rabbit polyclonal | 1:1000 | Sigma-Aldrich, St. Louis, MO, USA |
| $Na^+/K^+$-ATPase α (H-3) (sc-48345) | mouse monoclonal | 1:1000 | Santa Cruz Biotechnology, Dellas, TX, USA |
| anti-glyceraldehyde-3-phosphatase dehydrogenase (clone GAPDH-71.1) G8795 | mouse monoclonal | 1:2000 | Sigma-Aldrich, St. Louis, MO, USA |
| Anti-TurboGFP(d) (AB513) | rabbit polyclonal | 1:1000 | Evrogen, Moscow, Russia |
| HRP anti-rabbit (65-6120) | goat polyclonal | 1:10,000 | Thermo Fisher Scientific, Waltham, MA, USA |
| HRP anti-mouse | goat polyclonal | 1:10,000 | Invitrogen - Fisher Scientific, Waltham, MA, USA |
| HRP anti-guinea pig | goat polyclonal | 1:10,000 | Southern Biotec, Birmingham, AL, USA |
| HRP anti-rat (A9542) | rabbit polyclonal | 1:10,000 | Sigma-Aldrich, St. Louis, MO, USA |
